# An ancient polymorphic regulatory region within the BDNF gene associated with obesity modulates anxiety-like behaviour in mice and humans

**DOI:** 10.1101/2021.07.20.452916

**Authors:** Andrew R. McEwan, Benjamin Hing, Johanna C. Erickson, Yvonne Turnbull, Mirela Delibegovic, Felix Grassmann, Alasdair MacKenzie

## Abstract

Obesity and anxiety are morbidities notable for their increased impact on society during the recent COVID-19 pandemic. Understanding the mechanisms governing susceptibility to these conditions will increase quality of life and our resilience to future pandemics. In the current study we explored the function of a highly conserved regulatory region (BE5.1) within the BDNF gene that harbours a polymorphism strongly associated with obesity (rs10767664; p=4.69×10^−26^). Analysis in primary cells suggested that the major T-allele of BE5.1 was an enhancer whereas the obesity associated A-allele was not. However, CRISPR/CAS9 deletion of BE5.1 from the mouse genome (BE5.1KO) produced no significant effect on the expression of BDNF transcripts in the hypothalamus, no change in weight gain after 28 days and only a marginally significant increase in food intake. Nevertheless, transcripts were significantly increased in the amygdala of female mice and elevated zero maze and marble burying tests demonstrated a significant increase in anxiety-like behaviour that could be reversed by diazepam. Consistent with these observations, human GWAS cohort analysis demonstrated a significant association between rs10767664 and anxiousness in human populations. Intriguingly, interrogation of the human GTEx eQTL database demonstrated no effect on BDNF mRNA levels associated with rs10767664 but a highly significant effect on BDNF-antisense (BDNF-AS) gene expression and splicing suggesting a possible mechanism. We discuss our findings within the context of the known function and regulation of BDNF in obesity and anxiety whilst exploring the validity of interrogating GWAS data using comparative genomics and functional analysis using CRISPR genome editing in mice.

## Introduction

Since the first successful genome wide association study (GWAS) was carried out in a large affected human cohort (Ozaki et al., 2002) it has become clear that nearly all disease associated loci identified occur outside of gene coding regions (Pickrell, 2014) and within presumed regulatory regions (Maurano et al., 2012). Indeed, it was further posited that *“detailed mapping of cell-specific regulatory networks will be an essential task for fully understanding human disease biology”* (Boyle et al., 2017). However, this raises a major challenge because, compared to gene coding regions, the regulatory components that modulate these *cell-specific regulatory networks* are relatively unknown so their identification and characterisation remains difficult as does determining the effects of GWAS identified disease associated polymorphic variants on their activity.

The recent Covid-19 pandemic has highlighted the effects of obesity on Covid-19 mortality and the increased levels of anxiety suffered world-wide as a result of lockdown (Almandoz et al., 2020). Because of differing levels of susceptibility to the effects of Covid-19 in different populations a great deal of interest has focussed on the mechanisms regulating susceptibility to obesity and anxiety and the effects of genetic variation. The gene encoding the brain derived neurotrophic factor (BDNF) is particularly relevant in this regard because GWAS associated polymorphisms around the BDNF have been associated with conditions including depression (Hing et al., 2018), anxiety (Notaras and van den Buuse, 2020) and obesity(Speliotes et al., 2010; Thorleifsson et al., 2009). Of note is that the vast majority of disease associated SNPs in and around the BDNF gene locus are non-coding.

BDNF is a secreted protein that binds the TrkB receptor where it helps to support survival of existing neurons, encourages growth and differentiation of new neurons and synapses and is important in neuronal plasticity (Binder and Scharfman, 2004). The gene encoding the BDNF protein has 9 exons each with its own promoter (Pruunsild et al., 2007). Surprisingly, only exon 9 encodes the BDNF protein so the fact that the gene is controlled by 9 different promoters, that produce 17 different transcripts, speaks to the high level of transcriptional control required for proper cell-specific expression of the gene in the healthy brain (Pruunsild et al., 2007). Diverse BDNF transcripts, produced from different promoters, serve distinct molecular and behavioural functions including aggressiveness and modulation of components of the serotoninergic system that plays a role in mood modulation(Maynard et al., 2016) and appetite (Garfield and Heisler, 2009). In addition to promoter regions (Pruunsild et al., 2011), studies have also characterised many of the enhancer and repressor regions that support its tissue specific expression (Hing et al., 2012; Tuvikene et al., 2021). Efforts to identify other functional regulatory regions around the BDNF locus, such as enhancers, have employed high throughput next generation sequencing (NGS) technologies such as ATAC-seq (Song et al., 2019) and chromatin immunoprecipitation sequencing (ChIP-seq) to detect markers of active enhancers(H3K4me1, H3K27Ac) (Creyghton et al., 2010). These technologies have allowed the successful identification of an enhancer region (+3kb enhancer) within the rat BDNF intron 3 (Tuvikene et al., 2021). The same group were also successful in identifying a role for the CREB family of transcription factors in regulating BDNF and described an autoregulatory mechanism in BDNF regulation (Esvald et al., 2020).

Other efforts to identify BDNF regulatory regions combined genetic association analysis with comparative genomics to explore the effects of a polymorphism (rs12273363) associated with major depressive disorder(MDD) (Juhasz et al., 2011) on BDNF promoter activity. This study found that different alleles of rs12273363 changed the way that a highly conserved repressor region was able to filter the effects of different signal transduction pathways on BDNF promotor 4 activity (BP4) (Hing et al., 2012). It is therefore likely that combining both approaches; biochemical marker analysis and comparative genomics, with genome wide association analysis will be an effective combination to understand the tissue specific regulation of the BDNF gene and how its mis-regulation, due to allelic variation, may contribute to obesity and/or pathological anxiety.

In the current study we explored the functional biology of another SNP that had the fifth highest association to obesity (rs10767664; p=4.69×10^−26^) in a study of 204,158 individuals (Speliotes et al., 2010) and which also occurred in a highly conserved region (BE5.1) within intron 3 of the BDNF gene. Based on these observations, we hypothesised that this conserved region played a role in either appetite or metabolism. We, therefore, isolated BE5.1 from the human genome and reproduced both alleles of the rs10767664 polymorphism. We then analysed the effects of these allelic variants in reporter construct transformed into primary cell lines. CRISPR genome editing was then used to delete this region from the mouse genome and the effects of this deletion on BDNF gene expression and behaviour analysed. We finally conducted human cohort association analyses and eQTL analysis of rs10767664 allelic variants. The intriguing results of these experiments are discussed in the context of current efforts to derive functional data from GWAS association analysis.

## Materials and Methods

### Plasmid construction

Human BE5.1 was amplified using high fidelity PCR (Expand high fidelity system, Roche, UK) from human DNA (Cambio, UK) using the following primers; BE5.1 for. 5’-AATGAGGGAAAGTTTCACAGC-3’ and BE5.1 rev. 5’-CTGTGCCACTCTGCTCAAC-3’. Products were ligated into pGEM-T easy cloning vector (Promega, UK). All amplified DNA samples were sequenced (Source Bioscience, UK) to ensure correct sequence and orientation. To reproduce the A-variant the following primers were used for SDM by site directed mutagenesis using QuickChange II site directed mutagenesis kit (Agilent Technologies, UK) using the following primers; forward primer: 5’-GTAGGCTTGACATTGACATGTTTTTACTATTAATAATTTTAATTGGCTGAG-3’ and reverse primer 5’-CTCAGCCAATTAAAATTATTAATAGTAAAAACATGTCAATGTCAAGCCTAC -3’. BE5.1(A) and BE5.1(T) were then ligated into the BP4 luciferase reporter construct (based on pGL4-23) as previously described (Hing et al., 2012).

### Primary cell culture

Hypothalamic tissues were dissected from postnatal day 0-3 Sprague Dawley rat pups that were humanely sacrificed according to UK Home Office guidelines. Tissues were treated with 0.05% trypsin EDTA (Invitrogen, UK) for 15 mins at 37°C. Trypsin EDTA was replaced with soybean trypsin inhibitor (Sigma, UK) for 5 min at room temperature to stop reaction. This was then replaced with unsupplemented Neurobasal A (Invitrogen, UK) followed by mechanical dissociation. Cells were then resuspended in culture media [Neurobasal A, B27 (Invitrogen, UK), 1X glutamax (Invitrogen, UK) and Pen/strep (100 U/ml) (Invitrogen, UK)] and plated out at a density of ∼80000 viable cells/cm^2^on poly-L-lysine (20µg/ml) (Sigma, UK) pretreated plates following cell counting for viable cells using TC10™automated cell counter (Biorad) with trypan blue. Cells were incubated at 37ºC, 5% CO^2^ for 7 days prior to transfection.

### Transfections and treatments

All DNA constructs were quantified on a nanodrop machine (NanoDrop Technologies). Quantities of plasmid used for each transfection were adjusted to the size of each plasmid to ensure molar equivalence between experiments. Firefly luciferase plasmids were co-transfected with Renilla luciferase plasmid, pGL4.70 (Promega, UK), to normalize signals between transfections using magnetic particles (Neuromag;Oz Bioscience, UK) as described in manufacturer’s instructions. Cultures were incubated for 24 hrs prior to harvest for dual luciferase assay as per manufacturers instructions (Promega).

### Generation of gRNA molecules by a novel annealed oligo template (AOT) method

Single guide RNA (sgRNA) molecules were designed to disrupt the BE5.1 using the optimised CRISPR design tool (http://CRISPR.mit.edu/). sgRNA template was produced by annealing oligonucleotides to produce two different DNA templates using the annealed oligo template (AOT) method as previously described (Hay et al., 2019) that included a T7 polymerase binding site and predicted guide sequence target sites spanning the conserved region surrounding the BE5.1 enhancer (5’sgRNA; AACCCACTATAGCCCCCATGG and 3’sgRNA; TGGTTGCTGGCCCTCAGGTGG). These oligonucleotides were annealed and amplified using PCR to produce a 122 bp double strand sgRNA template. 100 ng of this template was used to produce sgRNA using a mMESSAGE mMACHINE T7 *in-vitro* transcription kit (Ambion) described in the manufacturer’s instructions and purified using a Megaclear kit (Ambion) with modifications as previously described (Harms et al., 2014).

### Production of genome edited mice

sgRNA molecules were microinjected at a concentration 10 ng/µl each into the cytoplasm of 1-cell C57/BL6 embryos as described (Harms et al., 2014) together with 10 ng/µl CAS9 mRNA (Life Technologies). Two-cell embryos were introduced into host CD1 mothers using oviduct transfer as previously described (Nagy, 2003) and correctly targeted offspring were determined by PCR of earclip DNA using the following flanking primers (AMX013; GCATCCTTTTGAGAGAAAT, AMX014; CCGAGGACACAGGAAGC).

### Quantitative reverse transcriptase-PCR

Brain tissues (hypothalamus and amygdala) were recovered from wild-type and BE5.1KO offspring and snap frozen on dry ice. Total RNA was extracted using the isolate II RNA minikit (Bioline). Quantitative reverse transcriptase-PCR (QrtPCR) on derived mouse cDNA was carried out using BDNF splice isoform specific primers (**See table S1**) to determine differential changes in BDNF splice forms using a Roche Light Cycler 480 with Roche SYBR green (Shanley et al., 2010; Shanley et al., 2011) normalised against the Non-POU Domain Containing Octamer Binding protein (*Nono)* housekeeping gene as previously described(Hay et al., 2020; Hay et al., 2019; McEwan et al., 2020a; McEwan et al., 2020b). .

### Animal studies

All animal studies were performed in accordance with UK Home Office guidelines and maintained under specific pathogen-free (SPF) facilities. The health status of these animals conformed to The Federation of European Laboratory Animal Science Associations (FELASA) guidelines whereby pathogen screening was carried out on a quarterly (3 monthly) basis using sentinel animals. BE5.1KO mice were maintained as a colony on a heterozygous C57BL/6 background which was mated to produce homozygous wild type and homozygous BE5.1KO age matched and sex matched individuals. These were identified using PCR of earclip DNA as described above. Once identified, animals were assigned random numbers to hide their genotype from the operators of subsequent tests. Male and female homozygous wildtype and BE5.1KO age matched littermates were housed under standard laboratory conditions (12 h light/12 h dark cycle), in plastic cages with food and water available ad libitum, depending on the experiment.

### Food intake studies

For food intake and high-fat diet preference studies, singly housed animals were provided with a choice of standard CHOW diet (low fat diet, LFD; 22.03 kcal% protein, 68.9kcal% carbohydrate and 9.08 kcal% fat) or high-fat diet (HFD; 20 kcal% protein, 20 kcal% carbohydrate and 60 kcal % fat; Research Diets Inc.) in different hoppers. The position of each hopper was changed regularly to rule out the possibility of position effects. Animals were weighed daily over a period of 23 days and LFD and HFD were also weighed daily to determine intake of each diet. Animals were also subjected to ECHO MRI analysis at the start and at the end of the feeding trial to assess changed in body fat mass v. total body mass.

### Elevated zero maze (EZM)

The EZM consists of an annular dark-gray platform (60 cm in diameter) constructed of opaque Perspex divided into four equal quadrants. Two opposite quadrants were “open”; the remaining two “closed” quadrants were surrounded by 16 cm high dark, opaque black walls. Quadrant lanes were 5 cm in width. Overhead lighting applying 100 lx at the level of the maze. The movement of animals was tracked using a camera and ANY-maze tracking software(Stoelting Europe). Distance travelled, average speed, number of crossings between light/dark zones and freezing episodes, freezing time and freezing latency were recorded.

### Marble burying test (MBT)

The MBT consisted of a a “EUROSTANDARD TYPE IV S” (Techiplast) cage (480 × 375 × 210 mm) cage with 5cm depth of wood chip bedding onto which 20 evenly spaced marbles were placed. Animals were then placed in the cage and the numbers of marbles buried after 30 minutes were recorded. This experiment was repeated with the same animals following i.p. injection of 10mg/kg of diazepam.

### Assessment of rs10767664 in human populations

We extracted the association of rs10767664 with all traits available in the MRC IEU OpenGWAS database [https://europepmc.org/article/ppr/ppr199104] with the function *phewas* from the *ieugwasr* package [Gibran Hemani (2021). ieugwasr: R interface to the IEU GWAS database API. R package version 0.1.5. URL: https://github.com/mrcieu/ieugwasr], as implemented in R [R Core Team (2020). R: A language and environment for statistical computing. R Foundation for Statistical Computing, Vienna, Austria. URL: https://www.R-project.org/.]. We restricted the results to all associations related to anxiety, worry, risk taking and sexual history. To account for multiple testing, we controlled the false-discovery rate to be less than 5% and thus report all associations with a Q-value of less than 5%. eQTL analysis of the effects of rs10767664 on human gene expression was undertaken by interrogation of the GTEx Portal (https://www.gtexportal.org/home/).

### Data analysis

From *in-vivo* pilot studies we calculated that a minimum of 6-12 animals per group would enable detection of a 25% difference between different parameters (anxiety-like behaviour, food intake) with 80% power using one-way ANOVA and/or general linear modelling. Statistical significance of data sets was analysed using either one-way analysis of variance (ANOVA) analysis with Bonferroni post hoc tests or using two tailed unpaired parametric Student *t*-test as indicated using GraphPad PRISM version 5.02 (GraphPad Software, La Jolla, CA, USA).

## Results

### The obesity associated polymorphism rs10767664 occurs within intron 3 of the BDNF gene and within a sequence highly conserved between humans, mice and amphibians

Because of its importance in obesity (Speliotes et al., 2010) and the importance of the BDNF gene in appetite control (Lebrun et al., 2006) we explored the hypothesis that allelic variation at the rs10767664 locus functionally changed the activity of an uncharacterised cis-regulatory element that controls aspects of the expression of the BDNF gene involved in appetite. We were intrigued to find that rs10767664 occurred within a 494 base pair region of conserved DNA (BE5.1; chr11:27,725,730-27,726,223) specifically within a 240bp region of the extreme conservation that demonstrated >90% conservation to amphibians; a depth of conservation spanning 360 million years of divergent evolution (Kumar and Hedges, 1998) **(Figure 1A and B)**. BE5.1 lay 2.4kb 5’ from promoter 4 of the BDNF gene (BP4) and within BDNF intron 3 (**Figure 1A**) and 20kb away from the recently reported +3kb enhancer (Tuvikene et al., 2021).

**Figure 1A.**
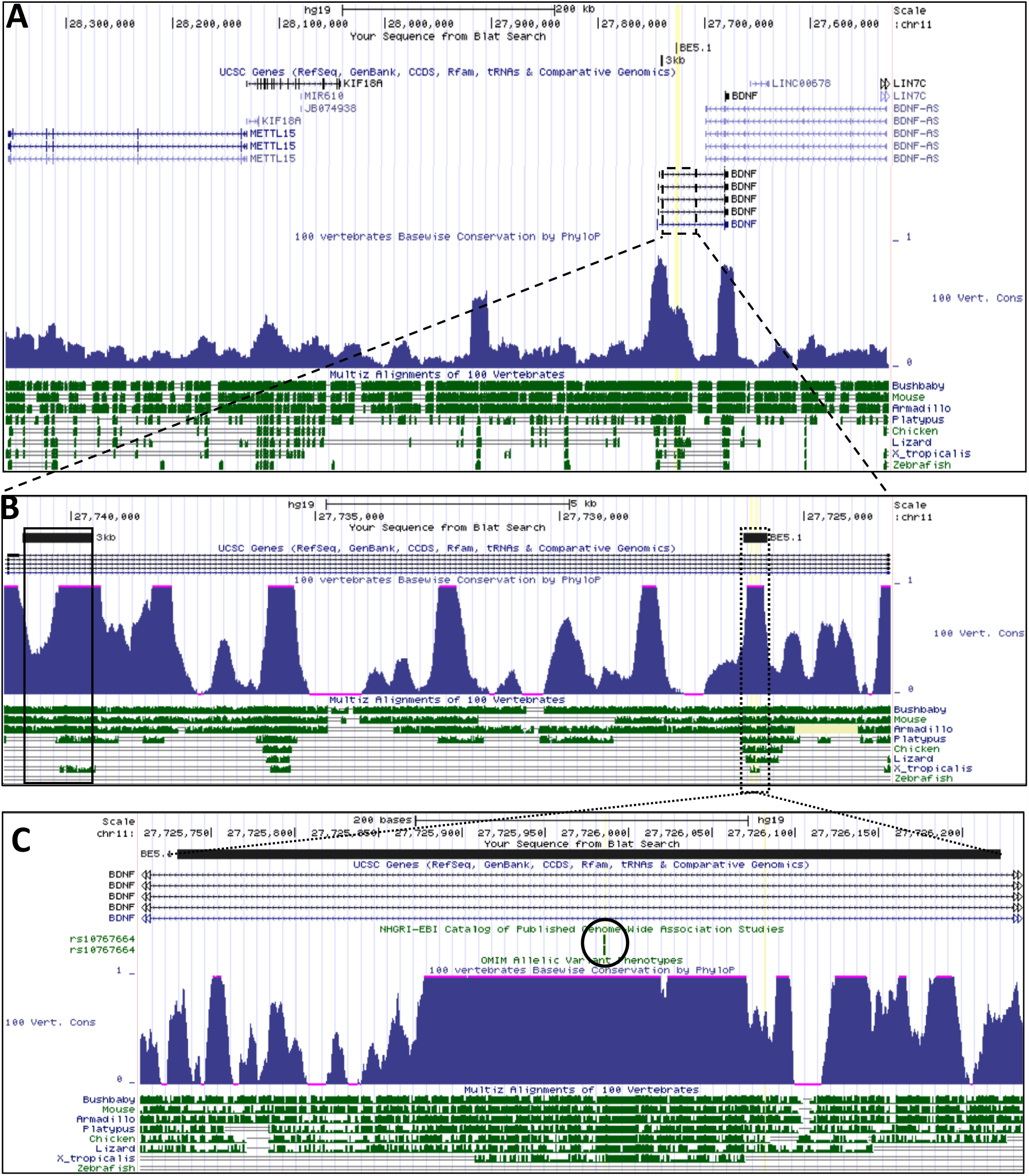
UCSC browser images covering 300kb of the human genome demonstrating the spatial relationships between human LIN7C, BDNF-AS, and BDNF genes. Intron 3, which harbours the BE5.1 regulatory region is highlighted by a dotted box. **(B)** Intron 3 of the BDNF gene representing the relationships between the BE5.1 enhancer explored in the current study, and the +3kb enhancer (3kb; Tuvikene et al., 2021) demonstrating relative conservation and depth of conservation through evolution (peaks and lines). **(C)** Demonstration of levels and depths of conservation of the BE5.1 regulatory region (X. tropicalis= 450 million years) that hosts the obesity associated rs10767664 polymorphism (circled in black).

### The T-allele of BE5.1 acts as an enhancer of BP4 activity in primary hypothalamic cells

To determine the effects of different alleles of BE5.1 on BDNF promoter activity we cloned 494 bp surrounding the human BE5.1 sequence using Hi-fidelity PCR and reproduced both human alleles using site directed mutagenesis. Both allelic variants were then cloned into a luciferase reporter plasmid containing the BP4 promoter region **(Figure 2)**, that represented the closest promoter to BE5.1, as previously described (Hing et al., 2012). We also chose this promoter as BP4 had previously been shown to be active in hypothalamus and amygdala in transgenic animals (Hing et al., 2012). These reporter plasmids were transfected into primary rat hypothalamic cells by magnetofection as previously described (Hing et al., 2012) and incubated for 48 hours. After this time cells were lysed and lysates analysed by dual luciferase assay. In this way we were able to demonstrate that the T-allele of BE5.1 could act as an enhancer of BP4 activity but that the obesity associated A-allele could not (**Figure 2**).

**Figure 2A.**
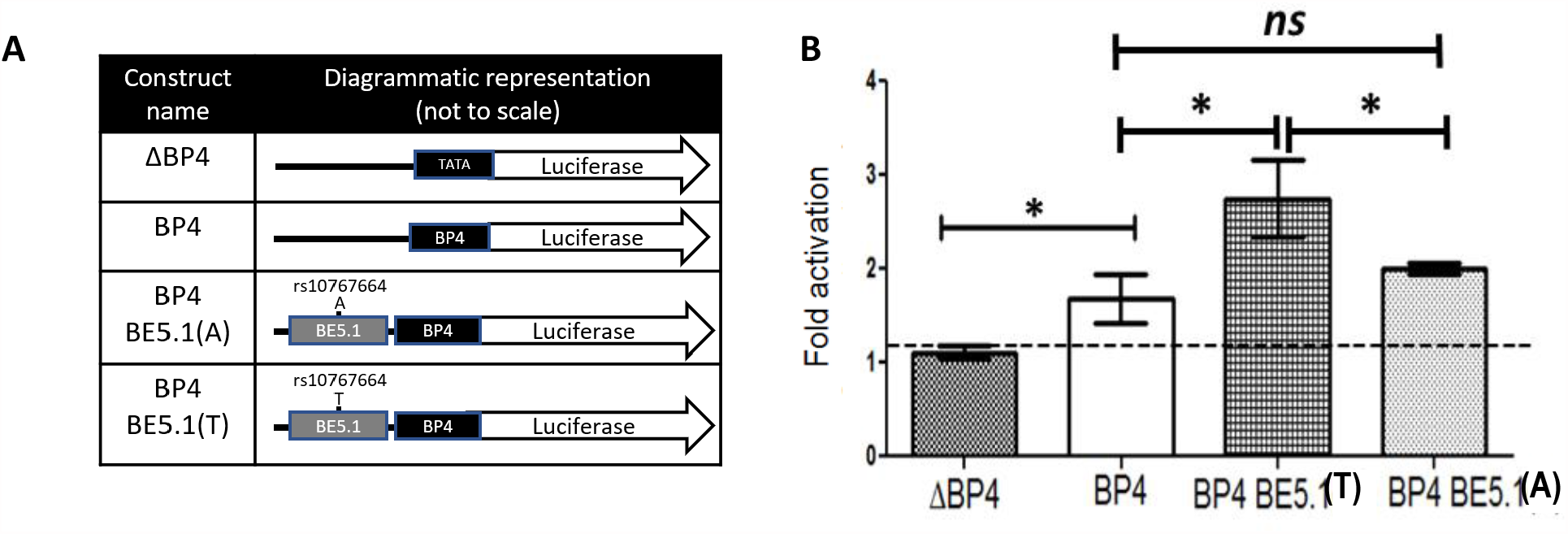
diagrammatic representation of the construct used in **2B**. Column 1; indicates the construct names used in **2B**. Column 2; a diagrammatic representation showing the relationship of each of the components. TATA; minimum TATA box promoter, BP4; BDNF promoter 4, BE5.1; BDNF enhancer 5.1. **Figure 2B** Bar graphs showing the firefly luciferase activities of each construct in **2A** 48 hours by dual luciferase analysis after magnetofection into primary hypothalamic cells (normalised to a co-transfected renilla luciferase constructs). Activity is expressed as fold activity compared to the minimal TATA box promoter construct (ΔBP4). *; p<0.05, *ns;* not significant, error bars; SEM.

### BE5.1 knockout animals are healthy and viable

We used CAS9/CRISPR genome editing to delete the BE5.1 enhancer from the mouse genome by cytoplasmic injection of guideRNA (gRNA) and CAS9 mRNA into 1-cell C57BL/6 mouse embryos as previously described (Hay et al., 2020; Hay et al., 2019; McEwan et al., 2020a; McEwan et al., 2020b). We were able to generate a single heterozygous female animal (BE5.1KO^+/-^) which was outbred on a C57BL/6 background for two generations to produce a colony of heterozygous male and female BE5.1 KO animals. Sequence analysis of the 5 most likely off target events for each gRNA was carried out as previously described (Hay et al., 2017) and could not detect any evidence of off-target changes within the genome of this line. These animals were then bred together to produce homozygous wild type (WT) and BE5.1KO animals, selected on the basis of PCR analysis, which proved to be healthy and viable.

### Deletion of BE5.1 has no significant effect on expression of BDNF isoforms in the hypothalamus

To determine the possible effects of the BE5.1 deletion on the expression of BDNF mRNA splice forms at a tissue specific level we recovered total RNA from the hypothalamus of male and female WT and BE5.1KO littermates. We then used quantitative reverse transcriptase PCR (QPCR) to quantify the expression of different isoforms of BDNF derived from different promoters **(Figure 3A)** in the hypothalamus where the expression of BDNF is known to influence appetite (Yeo and Heisler, 2012). However, we were unable to detect any significant difference in the expression of BDNF isoforms expressed from promoters 1-5 (exon 1-5) or promoter 9 (exon 9) in these samples **(Figure 3B-F)**. In order to determine the effects of deleting BE5.1 on the expression of the receptor for BDNF we also undertook QPCR analysis of the TrkB receptor but found no change in its expression **(Figure 3G)**.

**Figure 3A.**
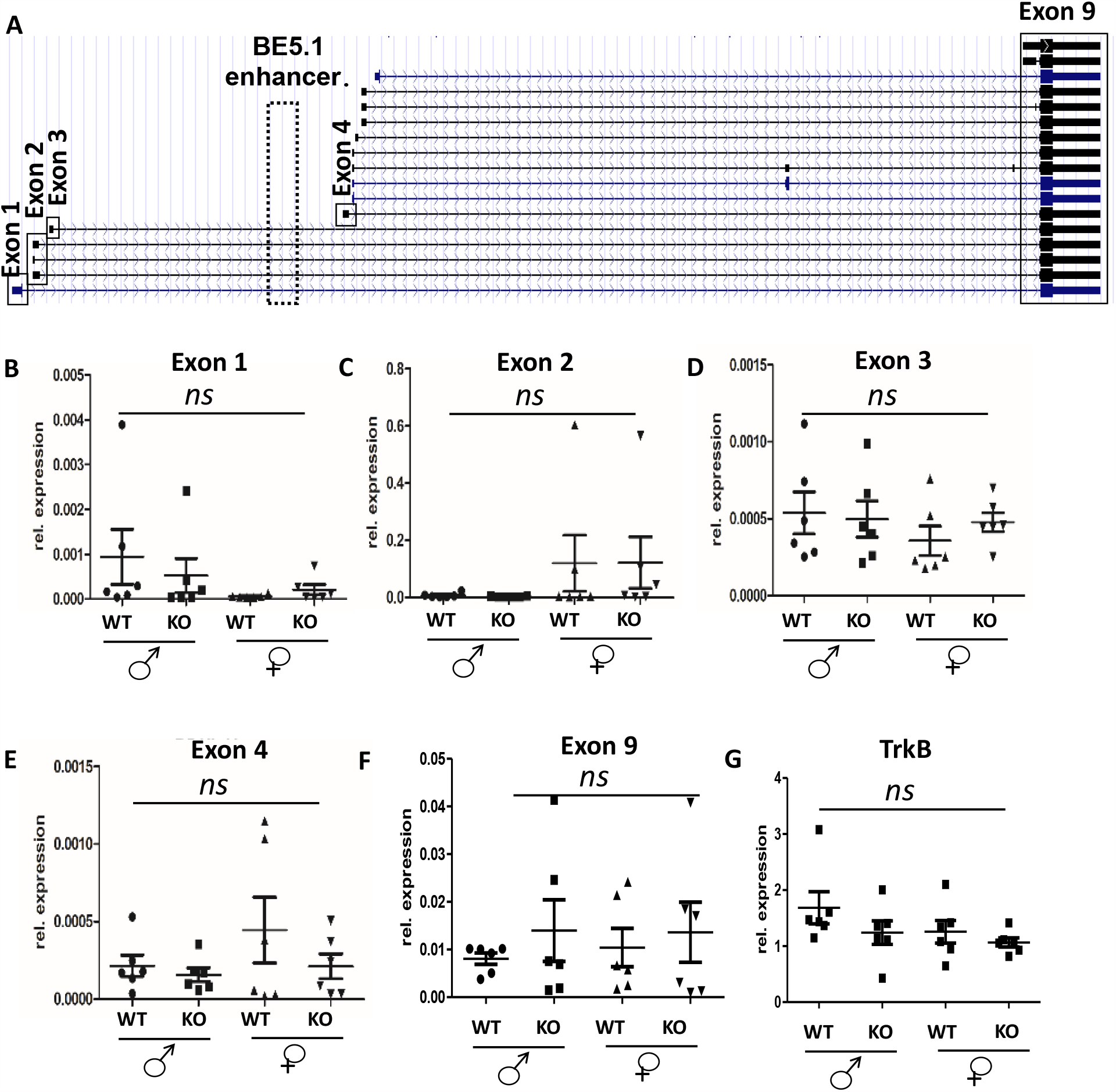
Figure derived from UCSC browser showing the different known BDNF mRNA isoforms and the spatial relationships between each promoter (black boxes) and the BE5.1 enhancer (broken box). **B-G**, results of QPCR analysis of BDNF and TRKB mRNA derived from the hypothalamus of wild type (WT) and BE5.1KO (KO) male and female mice comparing levels of BDNF mRNA isoform expression from promoters 1-4 and 9 (Exons 1-4 and 9) between WT and BE5.1KO male and female animals. *ns;* not significant, error bars; SEM.

### Deletion of the BE5.1 enhancer had no significant effect on weight gain and only marginal effects on food intake

Because the of the association of the rs10767664 polymorphism within BE5.1 with obesity, we explored the hypothesis that deletion of BE5.1 from the mouse genome would produce a significant change in food intake or weight gain. We allowed WT and BE5.1KO littermates access to a choice of either LFD diet or HFD for 28 days. Animals were weighed at regular intervals and analysed at the beginning and end of 28 days using echo MRI analysis. Although we detected a marginally significant increase in the intake of CHOW diet in BE5.1KO animals **(Figure 4 B and E)**, consistent with previously described effects of decreased in BDNF expression (Lebrun et al., 2006), we could not identify any significant differences in the intake of high fat diet, weight gain or fat distribution in these animals after 28 days suggesting that BE5.1 had only marginal effects on food intake and no significant effect on weight gain over the period of time studied **(Fig 4A, C, and D)**.

**Figure 4.**
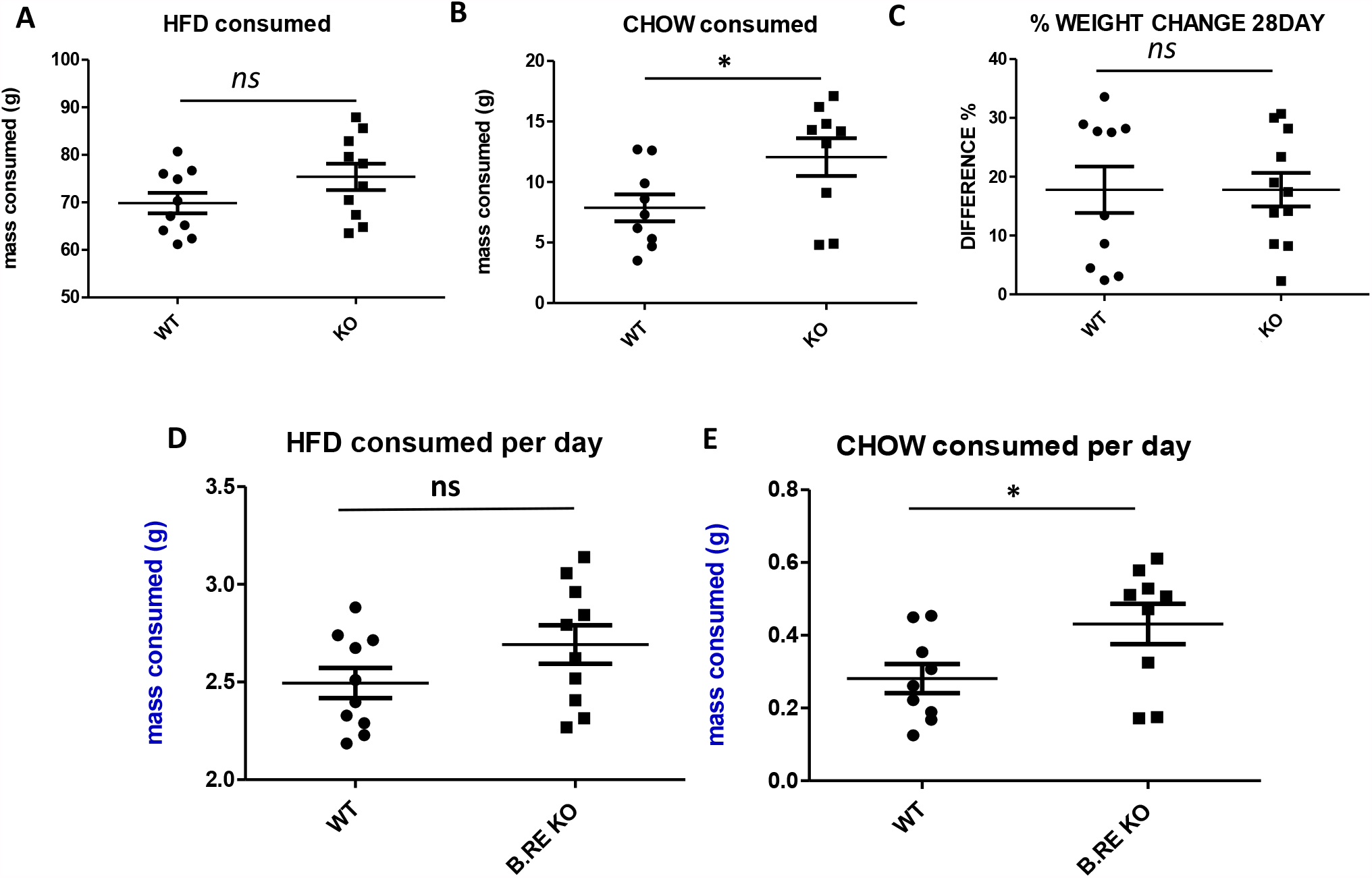
Scatter plots demonstrating the total mass of **(A)** High-fat diet and **(B)** Low-fat diet (CHOW) consumed either over the 28 days of the study (or **(D)** and **(E)** per day), comparing wild type (WT) and BE5.1KO (KO) animals. **C**. percentage change in total body mass comparing WT and KO animals exposed to a choice of high fat and CHOW diet over 28 days. n.s.; not significant, *; p<0.05.; error bars =SEM.

### Deletion of BE5.1 significantly up-regulates expression of BDNF exons in the female amygdala

In addition to the hypothalamus we also recovered tissues from the amygdala of male and female WT and BE5.1KO littermates. We used QPCR to quantify the expression of different exons of BDNF in the amygdala where the expression of BDNF is known to influence anxiety-like behaviour (Boyle, 2013). Although we observed a trend in the data towards increased expression of BDNF exons 4 and 9 in these samples in males the increase was only marginally significant for exon 4 (by t-test) and fell just short of significant for exon 9 (*p=0*.*051*) **(Figure 5A-E)**. However, compared to WT littermates we detected a significant increase in the expression of exons 1, 2 and 4, (excluding exon 3), and exon 9 in amygdala tissues derived from BE5.1KO female mice **(Figure 5A-E)**.

**Figure 5A-F.**
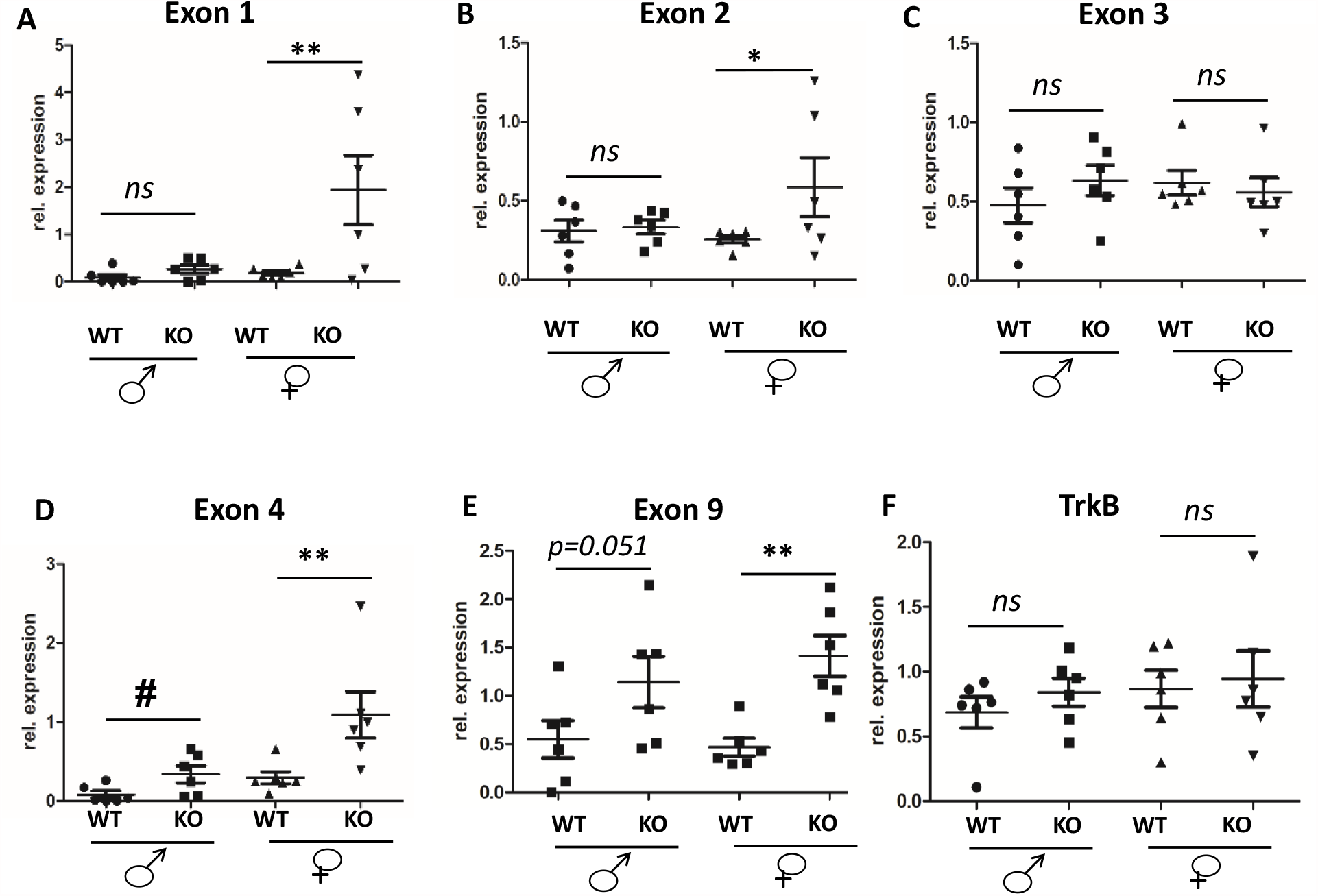
results of QPCR analysis of different exons of BDNF (A-E) and TRKB (F) mRNA derived from the amygdala region of wild type (WT) and BE5.1KO (KO) male and female mice comparing levels of mRNA expression between WT and BE5.1KO male and female animals normalised against Nono . n.s.; not significant, *; p<0.05. **; p<0.01 (by ANOVA); #; p<0.05 (two tailed t-test)..

### Female BE5.1KO mice demonstrate increased anxiety-like behaviour in the elevated zero maze

Because we detected increased levels of 4 different BDNF exons in the amygdala of female BE5.1KO mice we asked whether deletion of BE5.1 would influence anxiety-like behaviour in these animals. We subjected male and female BE5.1KO and WT littermates to the elevated zero maze (EZM) which is a robust method of detecting anxiety-like behaviour in mice (Heredia et al., 2013). However, we were unable to detect any significant difference between the time spent in the open quadrants **(Fig 6A)**, total distance travelled **(Fig 6B)**, Line crossings **(Fig 6C)** speed **(Fig6D)** or freezing time **(Fig 6E)** between male WT or BE5.1KO littermates in the EZM. By contrast, we observed highly significant changes in all of these behaviours in female BE5,1KO mice who spent significantly less time in the open quadrants of the EZM, showed reduced line crossings and overall speed with a highly significant increase in freezing time that accounted for much of the perceived lack of mobility in these animals **(Fig 6A-E)**.

**Figure 6A-E.**
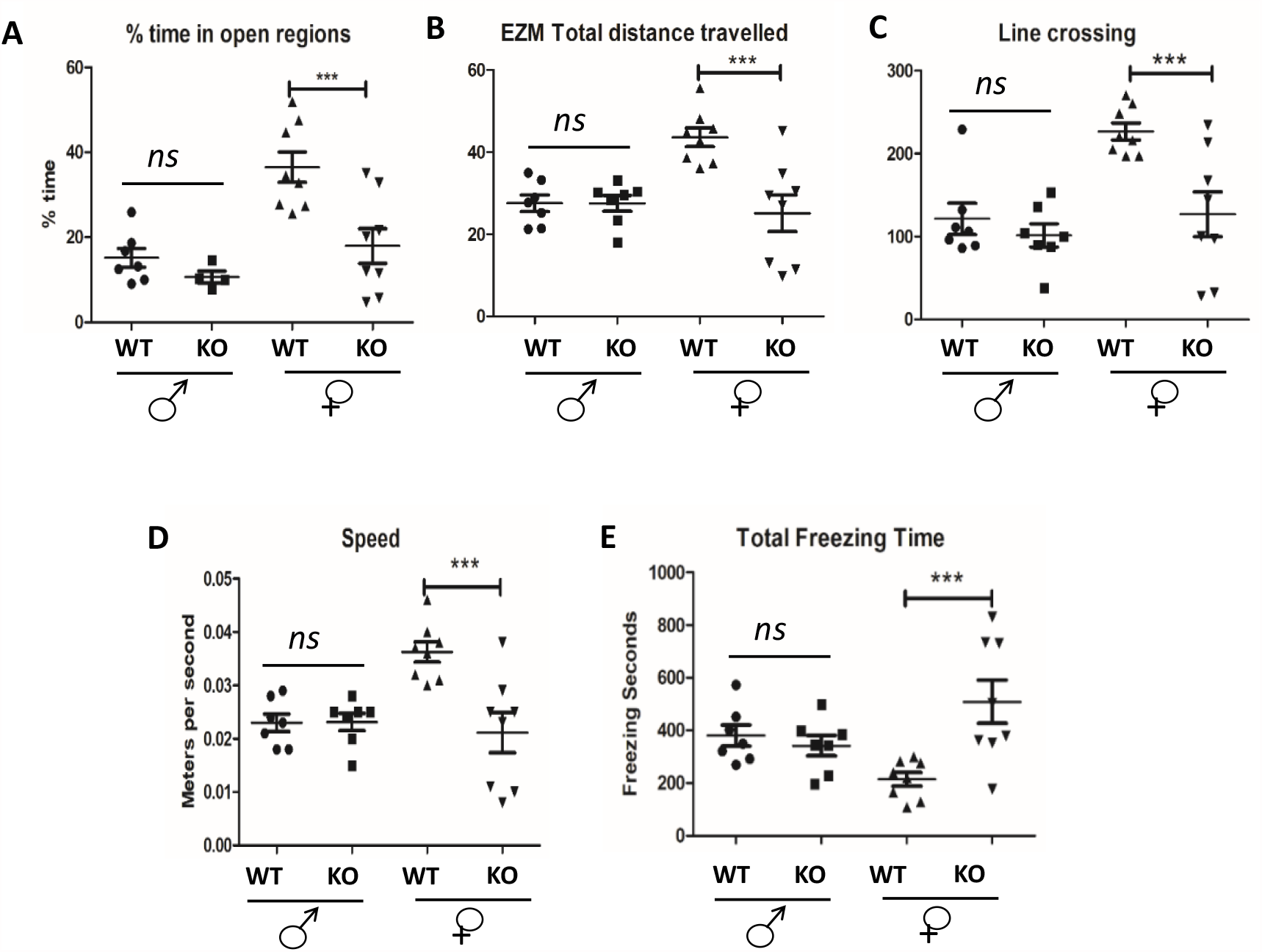
Scatterplots demonstrating the effects of deleting BE5.1 on the behaviour of wild type (WT) and BE5.1KO (KO) animals subjected to the elevated zero maze demonstrating (A) percentage time that each animal spent in the open quadrants of the maze, (B) the total distance travelled, (C) the number of crossings between the covered and open quadrants, (D) the average speed of each animal during the test and (E) the total time spent immobile (freezing behaviour) . ns; not significant, *; p<0.01. ***; p<0.005.

### Deletion of BE5.1 decreases marble burying in female mice which can be reversed using diazepam

In order to further test the effects of BE5.1 deletion on anxiety-like behaviour in mice we exposed male and female WT and BE5.1KO animals to a second anxiety test called the marble burying test (MBT); a well characterised test of anxiety-like behaviour in rodents (de Brouwer et al., 2019). Mice habitually bury marbles placed in their environment; a behaviour which decreases with increased anxiety. The advantage of the MBT is that, unlike the EZM; where animals become habituated to the test, the MBT can be repeated many times with the same animals with minimal change in the numbers of marbles each animal buries(de Brouwer et al., 2019). Briefly, the animal is placed in a cage with 3cm depth of fresh bedding (wood shavings) onto which 20 × 1.5 cm glass marbles are placed. After 30 minutes the animal is removed from the cage and the number of marbles buried are recorded. In the case of male animals, no significant difference was observed in the number of marbles buried by either the wild type or the BE5.1KO animals where both groups buried and average of 6-7 marbles each **(Fig 7A)**. However, consistent with our observations of the EZM, we detected a significant difference between the female WT and BE5.1KO whereby WT female mice buried an average of 5 marbles each whereas female BE5.1KO animals only buried between 1-2 marbles within the same time frame **(Fig 7A)**. Although marble burying has been widely used as a test of anxiety there are concerns that extrapolating anxiety-like behaviour from marble burying may be compromised by the parallel manifestation of obsessive-compulsive disorder related behaviours(de Brouwer et al., 2019). In order to better understand the contribution of anxiety-like behaviours in our results we undertook these tests again in females after treating a number of them with either a vehicle (saline) or saline containing 10mg/kg^-1^ of the anxiolytic drug diazepam. As before, female WT and BE5.1KO mice treated with the vehicle displayed a significant difference in the numbers of marbles buried within the 10 minutes of the study (**Fig 7B**). However, following treatment with diazepam, both groups of mice buried significantly more marbles and displayed no significant difference in the numbers of marbles buried (8-9; **Fig 7B**).

**Figure 7A.**
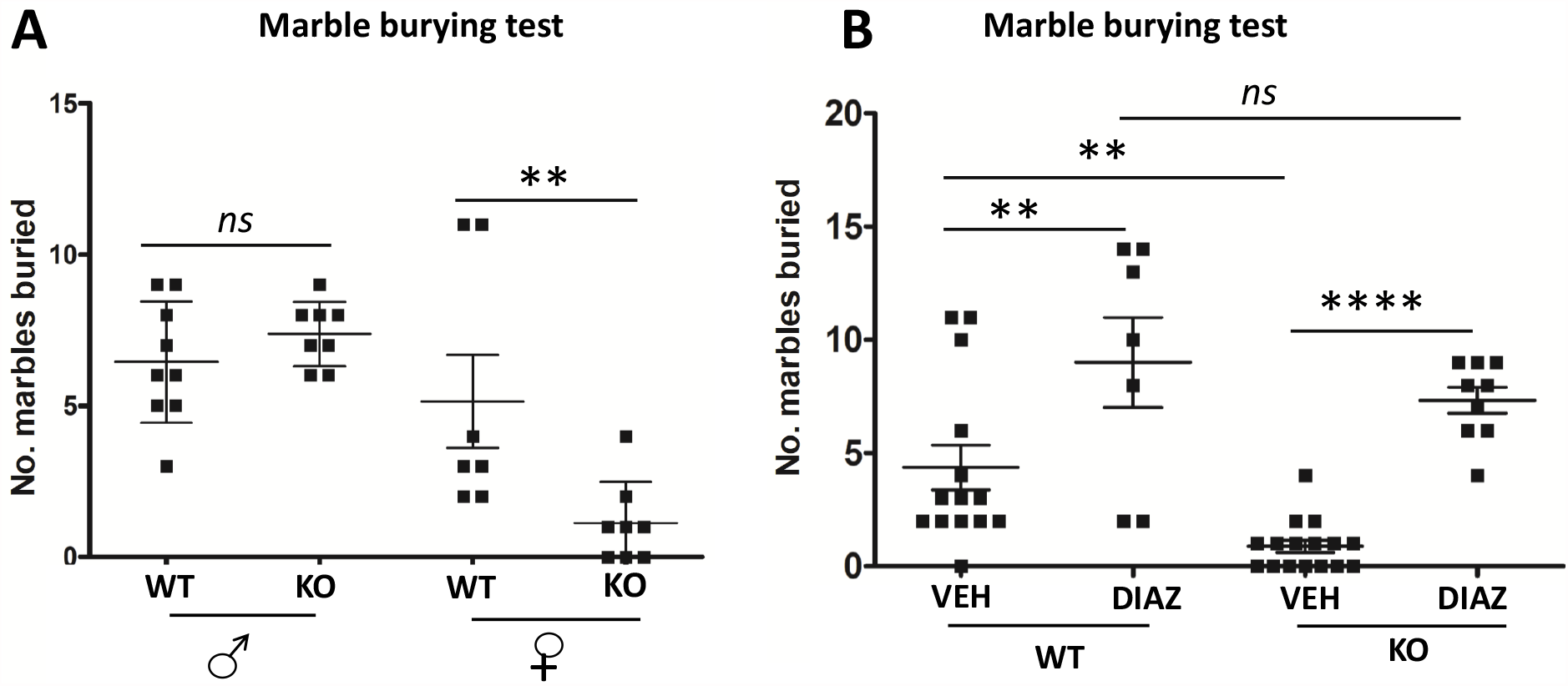
Scatterplots demonstrating the effects of deleting BE5.1 on marble burying behaviour in wild type (WT) and BE5.1KO (KO) animals. **B**. The effects of of diazepam (DIAZ; 10mg/kg^-1^ I.P.) on marble burying in female WT and BE5.1KO mice. *ns*; not significant, .**; p<0.01, ****; p<0.005, Error bars; SEM.

### The rs10767664 polymorphism is associated with feelings of anxiety and risk taking in human cohort studies

Next, we explored whether there was evidence for a functional role of rs10767664 in anxiety-like behaviour in humans. To this end, we extracted association results for rs10767664 from all genome-wide association studies (GWAS) included in the MRC IEU OpenGWAS database [https://europepmc.org/article/ppr/ppr199104]. We restricted the GWAS results to traits related to anxiety, worry, risk taking and sexual history. In total, we extracted 51 associations (**Supplementary Table S2**) of which 4 relating to anxiety/worry were statistically significant after adjustment for multiple testing and found that the A allele of rs10767664 was statistically significantly associated with anxious feelings and feelings of worry (FDR < 0.05, **Table 1**) consistent with the role for BE5.1 in anxiety consistent with our mouse data. In addition, the analysis also emphasised a high association with higher levels of overall risk taking, earlier age at first sexual intercourse and with a higher lifetime number of sexual partners (**Supplementary Table S2**).

**Table 1.** Statistically significant associations of the A allele of rs10767664 with traits related to human anxiety. *Beta: beta estimate for linear traits and log-odds ratio for binary traits, relative to the A allele of rs10767664, *SE: standard error of the estimate, *FDR: false-discovery rate.

| Trait | Trait ID | Sample Size | Beta* | SE** | P-value | FDR*** |
| --- | --- | --- | --- | --- | --- | --- |
| Worrier / anxious feelings | ukb-b-6519 | 450,765 | -0.005 | 0.001 | 3.30E-05 | 0.00056099 |
| Worrier / anxious feelings | ukb-a-51 | 497,161 | -0.006 | 0.001 | 9.14E-05 | 0.0009325 |
| Feeling worry | GCST006950 | 372,869 | -0.010 | 0.003 | 0.0002948 | 0.00250582 |
| Worry | GCST006478 | 348,219 | -0.010 | 0.003 | 0.0007492 | 0.00545849 |

### eQTL analysis of the rs10767664 polymorphism suggest a role for BE5.1 in regulating the expression of genes flanking the BDNF gene, including the BDNF-AS transcript, but not BDNF

The current study has demonstrated that the BE5.1 sequence acts as an enhancer in primary cells but that deleting BE5.1 from the mouse genome resulted in an increase in the expression of BDNF exonic mRNA in amygdala tissues in female mice. On the face of it, these observations appear to contradict each other as deleting an enhancer should not result in an increase in gene expression. For this reason, we asked whether different alleles of the rs10767664 polymorphism had an effect on the expression of genes surrounding BE5.1 (including the BDNF gene) in humans. We interrogated the GTEx database with the rs10767664 polymorphism and found a significant change in the expression of a number of genes flanking the BDNF locus including METTL15 (p=0.03 in amygdala) and LIN7C (strongly affected in vascular tissue (Aorta; p=4.5×10^−7^, tibial artery; p=1.2×10^−4^) and less strongly in esophagus (p=1.7×10^−4^), muscle (p=4.9×10^−3^) and visceral adipose tissue (p=0.006). In contrast, we were unable to find data supporting a change in the expression of the BDNF gene in GTEx as a result of the rs10767664 polymorphism. Intriguingly, however, rs10767664 had a significant effect on the expression of a gene called BDNF-AS; that expresses an alternatively spliced antisense RNA whose exon 5 sequence is complimentary to exon 9 of the BDNF mRNA transcript (Modarresi et al., 2012) **(Table 2)**. This observation is especially interesting as siRNA knockdown of BDNF-AS results in an increase in both BDNF mRNA and protein (Modarresi et al., 2012). The most significant effects of rs10767664 on BDNF-AS expression occurred in the nucleus accumbens; the addiction centre of the brain, and a number of cortical regions including the cingulate cortex that is a critical component of the limbic system and controls anxiety and fear (**Table 2**). We also observed that allelic variation at the rs10767664 polymorphism had significant effects on the splicing of the BDNF-AS primary transcript (**Table 2**).

**Table 2.** Effect of rs10767664 on gene expression and splicing of BDNF-AS in human brain tissue. p<0.05, highlighted in **bold**. NES=Normalized effect size, i.e. slope of the regression. From GTEx database.

|  |  | Single tissue eQTL |  | Single tissue splicing QTL |  |  |
| --- | --- | --- | --- | --- | --- | --- |
| Tissue | Samples | P-Value | NES | Intron Id [hg38] | P-Value | NES |
| Brain - Anterior cingulate cortex | 147 | **3.90E-03** | 0.17 | 27640005:27659171:clu_4862 | **7.80E-08** | -0.86 |
| Brain - Caudate | 194 | 0.08 | 0.11 | 27640005:27659171:clu_5504 | **2.10E-13** | -0.95 |
| Brain - Cerebellum | 209 | **0.02** | 0.18 | 27640005:27659171:clu_6041 | **1.60E-07** | -0.64 |
| Brain - Cortex | 205 | **8.80E-04** | 0.19 | 27640005:27659171:clu_5702 | **1.90E-09** | -0.75 |
| Brain - Frontal Cortex | 175 | **1.10E-04** | 0.21 | 27640005:27659171:clu_5288 | **6.20E-11** | -0.84 |
| Brain - Hippocampus | 165 | **0.01** | 0.13 | 27640005:27659171:clu_5002 | **3.20E-07** | -0.73 |
| Brain - Hypothalamus | 170 | 0.07 | 0.09 | 27640005:27659171:clu_5405 | **3.10E-07** | -0.73 |
| Brain - Nucleus accumbens | 202 | **9.70E-06** | 0.22 | 27640005:27659171:clu_5655 | **2.10E-14** | -1.00 |
| Brain - Putamen | 170 | **3.90E-03** | 0.17 | 27640005:27659171:clu_4853 | **3.90E-11** | -0.98 |
| Brain - Spinal cord | 126 | **1.70E-03** | 0.21 | 27640005:27659171:clu_4789 | **6.70E-10** | -0.95 |

## Discussion

The present study identified a region of deep conservation (350 million years), that we called BE5.1, within intron 3 of the BDNF gene that contained a human polymorphism highly associated with obesity (Speliotes et al., 2010). Magnetofection of primary hypothalamic cells, where BDNF has been shown to modulate appetite (Garfield and Heisler, 2009), demonstrated that the T-variant of BE5.1 acted as an enhancer of BDNF promoter 4 activity but that the obesity associated A-variant could not. In the context of the GWAS findings, this initial observation suggests that the T-allele of BE5.1, which exists within the majority of the population, promotes levels of BDNF gene expression in the hypothalamus appropriate for normal weight. These observations suggest that a contributing factor driving obesity in humans stems from a reduction in the ability of an enhancer element (BE5.1) to drive the expression of BDNF in the hypothalamus as a result of the obesity associated A-allele. This observation was consistent with previous studies showing that deletion of BDNF in the hypothalamus led to increased food intake (Lebrun et al., 2006).

Based on these observations, we used injection of CAS9 mRNA and sgRNA specific for the mouse BE5.1 enhancer to produce a line of mice deficient for BE5.1. Mice homozygous for the BE5.1 deletion were viable and appeared normal. Because the rs10767664 polymorphism had been strongly associated with obesity and was shown to occur within an enhancer of BP4 activity we expected to see a significant decrease of BDNF mRNA in hypothalamic tissues as well as an increase in food intake and greater weight gain in these animals. However, we did not detect any decrease in the mRNA expression of any of the five BDNF exons whose expression we analysed in the hypothalamus. Moreover, we saw no significant increase in HFD intake and no obvious change in weight gain over the 28 days of the feeding trial. Nevertheless, we did detect a marginally significant increase in CHOW diet intake suggesting that deleting BE5.1 had some effect on appetite. There are a number of possible explanations for these observations. First, it is possible that the role of the BE5.1 enhancer in appetite regulation does not reside in the hypothalamus but in a region of the brain, such as the nucleus accumbens (NAc), that regulates reward behaviour known to influence food intake(Cordeira et al., 2010). This hypothesis is supported by the observation that an allelic variant of the rs10767664 polymorphism, that is strongly associated with obesity in humans, is also associated with a highly significant change of the expression of the BDNF-AS gene in the NAc (GTEx eQTL analysis) to be further discussed later. Alternatively, because we analysed BDNF in the whole hypothalamus, it is possible that the regions of the hypothalamus where BDNF expression is influenced by BE5.1 may be too small to detect using QPCR. Finer dissection of the NAc, or different nuclei within the hypothalamus, followed by QPCR may be more revealing. In addition, we have yet to determine the tissue specificity of BE5.1 using reporter transgenic mice which we have done with other enhancers in the past (Davidson et al., 2011; Davidson et al., 2006; MacKenzie and Quinn, 1999; McEwan et al., 2020a; Miller et al., 2007; Miller et al., 2008; Shanley et al., 2010). Thus, although we were disappointed by the marginal nature of this result, we were satisfied that the small increase in CHOW diet intake was consistent with a possible role for BE5.1 in appetite and could form the basis of a larger study.

In contrast to our results in the hypothalamus, we found a significant increase in the expression of four BDNF mRNA isoforms in the amygdala of BE5.1KO female mice. Given the evidence above demonstrating that BE5.1 had enhancer like properties, the significant increase in BDNF expression in the amygdala of female BE5.1KO was not anticipated neither was the sex specificity of this effect. A further surprise was that deletion of BE5.1 was also associated with a highly significant increase in anxiety-like behaviour in these females in both the EZM and the MBT. The role of anxiety-like behaviour in our observations of the effects of BE5.1 deletion on the MBT was confirmed using the anxiolytic drug diazepam. Although the role of BDNF in the manifestation of mood disorders such as anxiety has been well known for some time, determining whether BDNF acts as an anxiolyic or anxiogenic regulator is far from resolved (Cazorla et al., 2011). For example, for many years it was accepted that BDNF or activation of the TrkB receptor reduced anxiety and depression (Martinowich et al., 2007). However, the results of other studies were consistent for a role for BDNF in increasing the pathophysiology of stress-induced anxiety (Berton et al., 2006; Eisch et al., 2003) and inhibition of the TrkB receptor has been actively explored as a potential anti-depressive treatment(Cazorla et al., 2011). Superficially, the coincidental observations of increased BDNF expression in amygdala and increased anxiety like behaviour in female mice would appear to support a role for increased BDNF expression in anxiety. However, we cannot, as yet, conclusively link one with the other and need to determine whether changes in the expression of the BDNF gene in the amygdala, or indeed some other gene in another part of the brain, are responsible. Again, this will be determined by closer dissection of the brains of BE5.1KO animals and the exploration of the effects of this deletion on the expression of other genes using next generation technologies such as RNA-seq. A further piece of evidence linking BE5.1 activity with anxiety came from our interrogation of the MRC IEU Open GWAS database. This analysis identified an association of the A-allele of rs10767664 with worry and anxious feelings in several large human cohorts. Again, direct functional comparisons of the effects of entirely deleting the BE5.1 enhancer against changing just 1 base pair within the enhancer are not strictly equivalent and we must temper our conclusions based on these differences. Nonetheless, if considered together; the increase in BDNF mRNA in the female amygdala as a result of deletion of BE5.1, together with the increase in anxiety like behaviour in female BE5.1KO mice, and the significant association of the A-allele of BE5.1 to anxiety in humans, provides a persuasive case for further studies of the role of BE5.1 in anxiety. Another intriguing observation from our analysis suggests a role for the A-allele in increased risk-taking behaviours and smoking further suggesting the involvement of BE5.1 in the NAc that controls addictive behaviours.

Our interrogation of the GTex database using rs10767664 also produced an unexpected result in that allelic variation of rs10767664 had no significant effect on the expression on the BDNF gene in any of the human tissues examined by GTEx. Instead, the GTEx data suggested that rs10767664 had a significant effect on the expression of the BDNF-AS gene. BDNF-AS does not encode a protein but produces a highly regulated and spliced mRNA whose transcriptional start site lies 200kb downstream from that of the BDNF gene and whose 5^th^ exon is complementary to exon 9 of the BDNF gene in both mice and humans(Modarresi et al., 2012). Modulation of gene expression at the transcriptional level by natural antisense (antisense encoded by the genome) has a number of different mechanisms including altering levels of DNA or histone methylation at gene promoters (Pelechano and Steinmetz, 2013). For example, siRNA knockdown of BDNF-AS resulted in an increase in both BDNF mRNA and a corresponding decrease in the repressive chromatin markers H3K27me3 at and EZH2 (component of the polycomb repressor complex; PRC2) at the BDNF promoter (Modarresi et al., 2012). It is therefore possible that a critical component of modulating levels of BDNF mRNA essential for modulating anxiety-like behaviour in the amygdala is governed by histone methylation and polycomb binding at the BDNF promoter region governed by BE5.1 driven BDNF-AS expression. Although a great deal of work remains to be done to determine the cogency of these hypotheses, the possibility that BE5.1 modulates mood through regulation of an antisense gene would be a unique finding with important repercussions for our understanding of the complexities of mood modulation and anxiety. We are currently exploring these possibilities using a combination of QPCR and RNA-seq of total RNA derived from BE5.1KO and WT amygdala and hypothalamus.

The discovery and analysis of two enhancer sequences within the BDNF intron 3 using different techniques provides an opportunity to compare the benefits and drawbacks of each approach. As mentioned in the introduction, previous attempts at locating enhancers controlling the BDNF gene were based on accepted indictors of enhancer activity such as open chromatin structure (DNAse-sensitivity), enrichment for specific chromatin markers such as H3K4me1, H3K4me3, and H3K27ac and CAGE analysis (bidirectional transcription) using cell culture based on line databases such as ENCODE and PHANTOM to identify a candidate enhancer that they called the +3kb enhancer (Tuvikene et al., 2021). This enhancer lay at the 5’ end of BDNF intron 3 roughly 20kb away from the BE5.1 enhancer that formed the basis of the current study. After its identification, Tuvikene et al (2021) used ChIP assays against the H3K27ac mark in mouse embryo forebrain cells and ATAC-seq in rat hippocampal and cortical cells to confirm its enhancer status. Further analysis of its effects on promoter 4 activity or expression of BDNF was determined by deleted the +3kb enhancer in cells using CRISPR genome editing. They subsequently identified many interacting transcription factors using an elegant combination of pull down and co-expression reporter studies in cortical neurones(Tuvikene et al., 2021). The authors also recognised that a significant fraction of the enhancer had been highly conserved in mammals (Tuvikene et al., 2021). Although this excellent study represented a state-of-the-art analysis of enhancer activity in cultured cells (dissociated primary cells and differentiated stem cells) the team were unable to describe the activity of the +3kb enhancer *in-vivo* or to determine its relevance to behaviour or human disease.

In parallel with Tuvikene et al (2021) we also sought enhancers that have an influence of the expression of the BDNF gene and used on-line data sources such as the UCSC browser that incorporates the latest GWAS data in combination with comparative analysis or over 100 vertebrates. Although we initially examined BE5.1 using chromatin markers available through ENCODE we concluded that the lack of these marks in BE5.1 reflected the inability of ENCODE cell lines to reproduce the specific contexts in which BE5.1 is active. Instead, our approach used a combination of evolutionary conservation and GWAS study association to identify a candidate enhancer element within BNDF with possible role in human disease susceptibility. We have demonstrated on many occasions that enhancers demonstrating high levels of tissue-specificity are often highly conserved through evolution(Davidson et al., 2011; Davidson et al., 2006; Hay et al., 2020; Hay et al., 2019; McEwan et al., 2021; McEwan et al., 2020a; Miller et al., 2007; Miller et al., 2008; Nicoll et al., 2012; Shanley et al., 2010). Furthermore, review of literature from other labs relating to well characterised enhancers also describe high degrees, and depth, of conservation (Lettice et al., 2017; Long et al., 2020; Oldridge et al., 2015; Tuvikene et al., 2021). Our initial functional analysis of BE5.1 was also similar to that employed by Tuvikene et al (2021) in that we found that the T-allele variant of BE5.1 could support activity of the BDNF promoter 4 but that the obesity associated A-variant could not. However, in contrast to Tuvikene et al (2021), we subsequently took advantage of CRISPR/CAS9 technology to delete the BE5.1 enhancer but in mice, as opposed to cell lines, and explored its function *in-vivo* where appropriate conditions for supporting BE.5.1 activity were present. In doing so we were also able to show that deletion of BE5.1 phenocopied aspects of the effects of the A-allele of rs10767664 in humans suggesting functional conservation. However, a disadvantage of this approach was that we were unable to carry out the extensive analysis of protein-DNA interactions achieved by Tuvikene et al (2021).

We conclude that both approaches demonstrate considerable merit and it is likely that combining them will be the key to, not only resolving the complex regulation of the BDNF locus, but also how these mechanisms are compromised by allelic variation and epigenetics.

## Acknowledgements

AMcE and AMcK were funded by a BBSRC project grant (BB/N017544/1) and Tenovus Scotland Grampian (award G19.08). We thank all the staff at the Medical Research Facility for their help and excellent advice in the completion of these studies. We also thank Giuseppe D’agostino for his guidance and help through the study.

## References

Almandoz, J.P., Xie, L., Schellinger, J.N., Mathew, M.S., Gazda, C., Ofori, A., Kukreja, S., and Messiah, S.Ed. (2020). Impact of COVID-19 stay-at-home orders on weight-related behaviours among patients with obesity. Clin Obes 10, e12386.

Berton, O., McClung, C.A., Dileone, R.J., Krishnan, V., Renthal, W., Russo, S.J., Graham, D., Tsankova, N.M., Bolanos, C.A., Rios, M., et al. . (2006). Essential role of BDNF in the mesolimbic dopamine pathway in social defeat stress. Science 311, 864–868.

Binder, D.K., and Scharfman, H.E. (2004). Brain-derived neurotrophic factor. Growth Factors 22, 123–131.

Boyle, E.A., Li, Y.I., and Pritchard, J.K. (2017). An Expanded View of Complex Traits: From Polygenic to Omnigenic. Cell 169, 1177–1186.

Boyle, L.M. (2013). A neuroplasticity hypothesis of chronic stress in the basolateral amygdala. Yale J Biol Med 86, 117–125.

Cazorla, M., Premont, J., Mann, A., Girard, N., Kellendonk, C., and Rognan, D. (2011). Identification of a low-molecular weight TrkB antagonist with anxiolytic and antidepressant activity in mice. J Clin Invest 121, 1846–1857.

Cordeira, J.W., Frank, L., Sena-Esteves, M., Pothos, E.N., and Rios, M. (2010). Brain-derived neurotrophic factor regulates hedonic feeding by acting on the mesolimbic dopamine system. J Neurosci 30, 2533–2541.

Creyghton, M.P., Cheng, A.W., Welstead, G.G., Kooistra, T., Carey, B.W., Steine, E.J., Hanna, J., Lodato, M.A., Frampton, G.M., Sharp, P.A., et al. . (2010). Histone H3K27ac separates active from poised enhancers and predicts developmental state. Proc Natl Acad Sci U S A 107, 21931–21936.

Davidson, S., Lear, M., Shanley, L., Hing, B., Baizan-Edge, A., Herwig, A., Quinn, J.P., Breen, G., McGuffin, P., Starkey, A., et al. . (2011). Differential activity by polymorphic variants of a remote enhancer that supports galanin expression in the hypothalamus and amygdala: implications for obesity, depression and alcoholism. Neuropsychopharmacology 36, 2211–2221.

Davidson, S., Miller, K.A., Dowell, A., Gildea, A., and Mackenzie, A. (2006). A remote and highly conserved enhancer supports amygdala specific expression of the gene encoding the anxiogenic neuropeptide substance-P. Mol Psychiatry 11, 323, 410-321.

de Brouwer, G., Fick, A., Harvey, B.H., and Wolmarans, W. (2019). A critical inquiry into marble-burying as a preclinical screening paradigm of relevance for anxiety and obsessive-compulsive disorder: Mapping the way forward. Cogn Affect Behav Neurosci 19, 1–39.

Eisch, A.J., Bolanos, C.A., de Wit, J., Simonak, R.D., Pudiak, C.M., Barrot, M., Verhaagen, J., and Nestler, E.J. (2003). Brain-derived neurotrophic factor in the ventral midbrain-nucleus accumbens pathway: a role in depression. Biol Psychiatry 54, 994–1005.

Esvald, E.E., Tuvikene, J., Sirp, A., Patil, S., Bramham, C.R., and Timmusk, T. (2020). CREB Family Transcription Factors Are Major Mediators of BDNF Transcriptional Autoregulation in Cortical Neurons. J Neurosci 40, 1405–1426.

Garfield, A.S., and Heisler, L.K. (2009). Pharmacological targeting of the serotonergic system for the treatment of obesity. J Physiol 587, 49–60.

Harms, D.W., Quadros, R.M., Seruggia, D., Ohtsuka, M., Takahashi, G., Montoliu, L., and Gurumurthy, C.B. (2014). Mouse Genome Editing Using the CRISPR/Cas System. Curr Protoc Hum Genet 83, 15 17 11–27.

Hay, E.A., Cowie, P., McEwan, A.R., Ross, R., Pertwee, R.G., and MacKenzie, A. (2020). Disease-associated polymorphisms within the conserved ECR1 enhancer differentially regulate the tissue-specific activity of the cannabinoid-1 receptor gene promoter; implications for cannabinoid pharmacogenetics. Hum Mutat 41, 291–298.

Hay, E.A., Khalaf, A.R., Marini, P., Brown, A., Heath, K., Sheppard, D., and MacKenzie, A. (2017). An analysis of possible off target effects following CAS9/CRISPR targeted deletions of neuropeptide gene enhancers from the mouse genome. Neuropeptides 64, 101–107.

Hay, E.A., McEwan, A., Wilson, D., Barrett, P., D’Agostino, G., Pertwee, R.G., and MacKenzie, A. (2019). Disruption of an enhancer associated with addictive behaviour within the cannabinoid receptor-1 gene suggests a possible role in alcohol intake, cannabinoid response and anxiety-related behaviour. Psychoneuroendocrinology 109, 104407.

Heredia, L., Torrente, M., Colomina, M.T., and Domingo, J.L. (2013). Assessing anxiety in C57BL/6J mice: a pharmacological characterization of the zero maze test. J Pharmacol Toxicol Methods 68, 275–283.

Hing, B., Davidson, S., Lear, M., Breen, G., Quinn, J., McGuffin, P., and MacKenzie, A. (2012). A polymorphism associated with depressive disorders differentially regulates brain derived neurotrophic factor promoter IV activity. Biol Psychiatry 71, 618–626.

Hing, B., Sathyaputri, L., and Potash, J.B. (2018). A comprehensive review of genetic and epigenetic mechanisms that regulate BDNF expression and function with relevance to major depressive disorder. Am J Med Genet B Neuropsychiatr Genet 177, 143–167.

Juhasz, G., Dunham, J.S., McKie, S., Thomas, E., Downey, D., Chase, D., Lloyd-Williams, K., Toth, Z.G., Platt, H., Mekli, K., et al. . (2011). The CREB1-BDNF-NTRK2 pathway in depression: multiple gene-cognition-environment interactions. Biol Psychiatry 69, 762–771.

Kumar, S., and Hedges, S.B. (1998). A molecular timescale for vertebrate evolution. Nature 392, 917–920.

Lebrun, B., Bariohay, B., Moyse, E., and Jean, A. (2006). Brain-derived neurotrophic factor (BDNF) and food intake regulation: a minireview. Auton Neurosci 126-127, 30–38.

Lettice, L.A., Devenney, P., De Angelis, C., and Hill, R.E. (2017). The Conserved Sonic Hedgehog Limb Enhancer Consists of Discrete Functional Elements that Regulate Precise Spatial Expression. Cell Rep 20, 1396–1408.

Long, H.K., Osterwalder, M., Welsh, I.C., Hansen, K., Davies, J.O.J., Liu, Y.E., Koska, M., Adams, A.T., Aho, R., Arora, N., et al. . (2020). Loss of Extreme Long-Range Enhancers in Human Neural Crest Drives a Craniofacial Disorder. Cell Stem Cell.

MacKenzie, A., and Quinn, J. (1999). A serotonin transporter gene intron 2 polymorphic region, correlated with affective disorders, has allele-dependent differential enhancer-like properties in the mouse embryo. Proc Natl Acad Sci U S A 96, 15251–15255.

Martinowich, K., Manji, H., and Lu, B. (2007). New insights into BDNF function in depression and anxiety. Nat Neurosci 10, 1089–1093.

Maurano, M.T., Humbert, R., Rynes, E., Thurman, R.E., Haugen, E., Wang, H., Reynolds, A.P., Sandstrom, R., Qu, H., Brody, J., et al. . (2012). Systematic localization of common disease-associated variation in regulatory DNA. Science 337, 1190–1195.

Maynard, K.R., Hill, J.L., Calcaterra, N.E., Palko, M.E., Kardian, A., Paredes, D., Sukumar, M., Adler, B.D., Jimenez, D.V., Schloesser, R.J., et al. . (2016). Functional Role of BDNF Production from Unique Promoters in Aggression and Serotonin Signaling. Neuropsychopharmacology 41, 1943–1955.

McEwan, A., Erickson, J.C., Davidson, C., Heijkoop, J., Turnbull, Y., Delibegovic, M., Murgatroyd, C., and MacKenzie, A. (2021). The anxiety and ethanol intake controlling GAL5.1 enhancer is epigenetically modulated by, and controls preference for, high-fat diet. Cell Mol Life Sci 78, 3045–3055.

McEwan, A.R., Davidson, C., Hay, E., Turnbull, Y., Erickson, J.C., Marini, P., Wilson, D., McIntosh, A.M., Adams, M.J., Murgatroyd, C., et al. (2020a). CRISPR disruption and UK Biobank analysis of a highly conserved polymorphic enhancer suggests a role in male anxiety and ethanol intake. Mol Psychiatry.

McEwan, A.R., Erickson, J.C., Davidson, C., Heijkoop, J., Turnbull, Y., Delibegovic, M., Murgatroyd, C., and MacKenzie, A. (2020b). The anxiety and ethanol intake controlling GAL5.1 enhancer is epigenetically modulated by, and controls preference for, high fat diet. Cellular and Molecular Life Sciences In Press, In Press.

Miller, K.A., Barrow, J., Collinson, J.M., Davidson, S., Lear, M., Hill, R.E., and Mackenzie, A. (2007). A highly conserved Wnt-dependent TCF4 binding site within the proximal enhancer of the anti-myogenic Msx1 gene supports expression within Pax3-expressing limb bud muscle precursor cells. Dev Biol 311, 665–678.

Miller, K.A., Davidson, S., Liaros, A., Barrow, J., Lear, M., Heine, D., Hoppler, S., and MacKenzie, A. (2008). Prediction and characterisation of a highly conserved, remote and cAMP responsive enhancer that regulates Msx1 gene expression in cardiac neural crest and outflow tract. Dev Biol 317, 686–694.

Modarresi, F., Faghihi, M.A., Lopez-Toledano, M.A., Fatemi, R.P., Magistri, M., Brothers, S.P., van der Brug, M.P., and Wahlestedt, C. (2012). Inhibition of natural antisense transcripts in vivo results in gene-specific transcriptional upregulation. Nat Biotechnol 30, 453–459.

Nagy, A.G., M. Vinterstein, K and Behringer, R. (2003). Manipulating the Mouse Embryo, third edition edn (Cold Spring Harbor: Cold Spring Harbor laboratory Press).

Nicoll, G., Davidson, S., Shanley, L., Hing, B., Lear, M., McGuffin, P., Ross, R., and MacKenzie, A. (2012). Allele-specific differences in activity of a novel cannabinoid receptor 1 (CNR1) gene intronic enhancer in hypothalamus, dorsal root ganglia, and hippocampus. J Biol Chem 287, 12828–12834.

Notaras, M., and van den Buuse, M. (2020). Neurobiology of BDNF in fear memory, sensitivity to stress, and stress-related disorders. Mol Psychiatry 25, 2251–2274.

Oldridge, D.A., Wood, A.C., Weichert-Leahey, N., Crimmins, I., Sussman, R., Winter, C., McDaniel, L.D., Diamond, M., Hart, L.S., Zhu, S., et al. . (2015). Genetic predisposition to neuroblastoma mediated by a LMO1 super-enhancer polymorphism. Nature 528, 418–421.

Ozaki, K., Ohnishi, Y., Iida, A., Sekine, A., Yamada, R., Tsunoda, T., Sato, H., Sato, H., Hori, M., Nakamura, Y., et al. . (2002). Functional SNPs in the lymphotoxin-alpha gene that are associated with susceptibility to myocardial infarction. Nat Genet 32, 650–654.

Pelechano, V., and Steinmetz, L.M. (2013). Gene regulation by antisense transcription. Nat Rev Genet 14, 880–893.

Pickrell, J.K. (2014). Joint analysis of functional genomic data and genome-wide association studies of 18 human traits. Am J Hum Genet 94, 559–573.

Pruunsild, P., Kazantseva, A., Aid, T., Palm, K., and Timmusk, T. (2007). Dissecting the human BDNF locus: bidirectional transcription, complex splicing, and multiple promoters. Genomics 90, 397–406.

Pruunsild, P., Sepp, M., Orav, E., Koppel, I., and Timmusk, T. (2011). Identification of cis-elements and transcription factors regulating neuronal activity-dependent transcription of human BDNF gene. J Neurosci 31, 3295–3308.

Shanley, L., Davidson, S., Lear, M., Thotakura, A.K., McEwan, I.J., Ross, R.A., and MacKenzie, A. (2010). Long-range regulatory synergy is required to allow control of the TAC1 locus by MEK/ERK signalling in sensory neurones. Neurosignals 18, 173–185.

Shanley, L., Lear, M., Davidson, S., Ross, R., and MacKenzie, A. (2011). Evidence for regulatory diversity and auto-regulation at the TAC1 locus in sensory neurones. J Neuroinflammation 8, 10.

Song, M., Yang, X., Ren, X., Maliskova, L., Li, B., Jones, I.R., Wang, C., Jacob, F., Wu, K., Traglia, M., et al. . (2019). Mapping cis-regulatory chromatin contacts in neural cells links neuropsychiatric disorder risk variants to target genes. Nat Genet 51, 1252–1262.

Speliotes, E.K., Willer, C.J., Berndt, S.I., Monda, K.L., Thorleifsson, G., Jackson, A.U., Lango Allen, H., Lindgren, C.M., Luan, J., Magi, R., et al. . (2010). Association analyses of 249,796 individuals reveal 18 new loci associated with body mass index. Nat Genet 42, 937–948.

Thorleifsson, G., Walters, G.B., Gudbjartsson, D.F., Steinthorsdottir, V., Sulem, P., Helgadottir, A., Styrkarsdottir, U., Gretarsdottir, S., Thorlacius, S., Jonsdottir, I., et al. . (2009). Genome-wide association yields new sequence variants at seven loci that associate with measures of obesity. Nat Genet 41, 18–24.

Tuvikene, J., Esvald, E.E., Rahni, A., Uustalu, K., Zhuravskaya, A., Avarlaid, A., Makeyev, E.V., and Timmusk, T. (2021). Intronic enhancer region governs transcript-specific Bdnf expression in rodent neurons. Elife 10.

Yeo, G.S., and Heisler, L.K. (2012). Unraveling the brain regulation of appetite: lessons from genetics. Nat Neurosci 15, 1343–1349.

