## Supplementary Table 1 for "An ancient polymorphic regulatory region within the BDNF gene associated with obesity modulates anxiety-like behaviour in mice and humans"

| Primer name | Sequence (5’-3’) |
|---|---|
| mBDNFI | GTGTGACCTGAGCAGTGGGCAAAGGA |
| mBDNFII | GGAAGTGGAAGAAACCGTCTAGAGCA |
| mBDNFIII | GCTTTCTATCATCCCTCCCCGAGAGT |
| mBDNFIV | CTCTGCCTAGATCAAATGGAGCTTC |
| mBDNFIXA | CCCAAAGCTGCTAAAGCGGGAGGAAG |
| mBDNFrev: | GAAGTGTACAAGTCCGCGTCCTTA |
| TRKBFor | CTGGGGCTTATGCCTGCTG |
| TRKBRev: | AGGCTCAGTACACCAAATCCTA |

**Supplementary Table 1.** QPCR primers used to determine the effects of deleting BE5.1 on the expression of different isoforms of BDNF.
